# Establishment of chicken muscle and adipogenic cell cultures for cultivated meat production

**DOI:** 10.1101/2025.05.09.652510

**Authors:** Vanessa Haach, Karine R. D. Silveira, Maíra de A. Peixoto, Ana Paula P. Sá, Vanessa Gressler, Vivian Feddern, Adriana M. G. Ibelli, Luciano Paulino Silva, Ana Paula Bastos

## Abstract

Cultured meat aims to replicate the sensory and functional properties of conventional meat by developing structured muscle tissue using cell culture. This study focuses on the culture of chicken embryonic and mesenchymal stem cells (MSCs) to derive muscle, and fat, optimizing conditions for differentiation and integration. We utilized monolayer and three-dimensional microcarrier-based cultures to produce muscle fibers and adipocytes while maintaining the extracellular matrix (ECM) integrity essential for tissue cohesion. Key pluripotency and myogenic markers (e.g., *cOCT4*, *cMYOD*, *cMYH1E*) were analyzed during differentiation, revealing dynamic gene expression patterns that underscore myogenesis. Myoblast differentiation into mature myotubes demonstrated decreased *cPAX7* and increased *cMYMK*, confirming lineage commitment and muscle fiber formation. Adipogenesis was induced in MSCs using food-grade lecithin, which activated PPARγ, C/EBPα, and FABP4, resulting in robust lipid droplet accumulation. To scale production, microcarriers facilitated cell proliferation, while transglutaminase-based stabilization enabled the formation of three-dimensional tissue structures comparable to native meat. Our findings highlight advances in culture protocols, genotypic and phenotypic expression analyses of multinucleated chicken muscle and adipocyte cells for cultured meat production.

## 1. Introduction

Over the past decade, the market has seen major shifts in consumer demand and product innovation for meat alternatives. Consequently, it is of paramount importance to sustain research efforts and develop alternative methods and proteins for the generation of novel products. Cellular agriculture is a new sector that attempts to provide a more sustainable alternative to traditional animal product production, eliminating the need for animal slaughter. Among the most promising developments in meat alternatives is the production of animal proteins from animal cell cultures or cultured meat. These advancements have the capacity to tackle various pressing challenges such as scarcity of food, climate change, worries regarding animal well-being, and difficulties related to public health [1–3]. Nonetheless, this technique faces numerous technological obstacles.

The primary ingredient in cultured meat is animal cell lines [4]. An indispensable requirement for any bioprocess, particularly in the successful creation of cultured meat, is the presence of a cell line that exhibits consistent and replicable features. Nevertheless, the absence of availability to thoroughly characterized cell lines poses a substantial obstacle to the investigation of cultured meat. It is essential that the initial cell types exhibit a high proliferation rate or self-renewal capacity in order to attain sufficient quantities for the effective production of cultured meat. Additionally, these cells must be capable of differentiating into the completely developed cell types that constitute meat [5].

There are two notable strategies for establishing cell lines for cultured meat production: (1) utilizing a sample of the tissue of interest (primary cell sources), coupled with the isolation of progenitor cells residing in the muscle; and (2) employing pluripotent or multipotent stem cell sources, such as embryonic stem cells (ESCs) and induced pluripotent stem cells (iPSCs), which possess the capacity to differentiate into muscle-resident progenitor cells [6]. Although stem cells, such as muscle stem cells and pluripotent stem cells, are widely employed as a cellular source for cultured meat, they are uncommon in the animal body and difficult to multiply on a large scale. Conversely, somatic cells, which compose the majority of the body, can be effectively transdifferentiated into muscle cells under specific circumstances.

The main cellular constituents of meat include skeletal myocytes, adipocytes, fibroblasts, chondrocytes, and hematopoietic cell types [7]. To optimize the production of cultured meat, it is crucial to determine the specific cultivation conditions that promote muscle cell proliferation and differentiation of satellite cells into myotubes and myofibers. These conditions should also preserve meat-like texture and flavor characteristics [8]. Supplementary methodologies are required to isolate target cells and achieve additional purification from these preliminary cell extracts. Consequently, a variety of techniques are employed to purify cells, each of which possesses a distinct set of benefits and drawbacks. Current cell separation methods utilize surface proteins, differential adhesion, selective plating, genetic expression, and cell detachment [4]. In addition, physical principles are employed to effectively isolate specific cells based on their phenotypic characteristics, including cell sorting (FACS) and capture using magnetic beads that are coated with cell-specific antibodies [9,10].

The viability of the cultured meat sector largely depends on technological advancements in both industry and research. The majority of research conducted on cultured meat has mostly been on producing a product that mimics the appearance and texture of fresh meat by proliferating and differentiating muscle stem cells [11–13]. However, producing sufficient samples of cultured meat for sensory panel testing remains a challenge [13], which makes it difficult to evaluate the technical flavor and texture attributes. Few studies have demonstrated evidence about the nutritional composition of cell lines, observing them as ingredients. In spite of these obstacles, it is possible to investigate and enhance muscle satellite cell culture techniques to guarantee that they exhibit flavor characteristics that are comparable to those of conventional meat. Our study was designed to resolve technological deficiencies and reinforce initiatives within the cultured meat industry. Our initiative was dedicated to the development of essential ingredients for the production of cultivated food products, specifically muscular and adipocyte chicken cell lines.

## 2. Materials and Methods

### 2.1. Chicken embryonic stem cells (blastoderm) isolation

The use of animals was approved by the Animal Use Ethics Committee of Embrapa Suínos e Aves (protocol number 22/2022). All experimental processes were conducted in accordance with the standard procedure. Blastoderm cells at stage X of Eyal-Giladi and Kochav (EGK) [14] were isolated from unincubated fertile eggs of specific pathogen-free (SPF) chickens 20–23 hours after fertilization. A piece of filter paper with a central aperture was placed gently onto the vitelline membranes, in order to frame the blastoderm. Afterwards, the vitelline membranes around the filter paper were cut, washed with Dulbecco’s phosphate-buffered saline (DPBS; Thermo Scientific) containing 1% antibiotic-antimycotic (Thermo Scientific) to remove the yolk. The cells were centrifuged and then filtered through 100, 70 e 40 μm strainer (Corning). The cells were collected by centrifugation and resuspended in Dulbecco’s modified Eagle’s medium (DMEM) low glucose (Thermo Scientific) supplemented with 10% fetal bovine serum (FBS; Thermo Scientific) and 1% antibiotic-antimycotic (Thermo Scientific).

### 2.2. Chicken mesenchymal stem cells and muscle satellite cells isolation

Embryonated SPF chicken eggs were incubated at 37.5 °C and 55% humidity for 15 days and selected by ovoscopy. Mesenchymal stem cells (MSCs) and satellite muscle cells were isolated from SPF chicken embryos within 15 days of incubation. Muscles from the thorax and hind limbs were collected and washed with DPBS (Thermo Scientific) containing 1% antibiotic-antimycotic (Thermo Scientific). The muscle tissue was cut into small fragments using scissors on a glass plate. The minced tissue was dissociated using 0.1% collagenase type I (Thermo Scientific), incubated at 37 °C for 1 hour, and during digestion, gently triturated by suction in a syringe with an 18-gauge needle, 10 times every 15 minutes, and after that, centrifuged. Subsequently, the digestion tissue was incubated with 0.25% trypsin-EDTA (Thermo Scientific) at 37 °C for 20 minutes, and then FBS (Thermo Scientific) was added to neutralize trypsin, and centrifuged. The cell suspension was filtered through 100, 70, and 40 μm strainer (Corning), and centrifuged. Afterward, the red blood cells were lysed using Pharm Lyse™ Buffer (BD Biosciences), incubated for 10 minutes at 4 °C, added to DMEM medium, and centrifuged. The cells were cultured in a grown medium composed of DMEM high glucose medium (Thermo Scientific) supplemented with 20% FBS (Thermo Scientific) and 1% antibiotic-antimycotic (Thermo Scientific), at 37 °C under 5% CO_2_. The cells were plated in T75 flasks, and after 2 hours the adherent cells were obtained and identified as MSCs. The supernatant containing non-adherent cells was collected to proceed with the selective adhesion of chicken muscle satellite cells.

### 2.3. Chicken muscle satellite cell culture and cell differentiation

The muscle satellite cells were selected by selective adhesion. The collected supernatant was plated in new T75 flasks and cultured for 1 day. This supernatant was transferred to new T75 flasks and cultured for another day. The following day, adherent cells were detached with 0.05% trypsin-EDTA for 5 minutes, centrifuged and resuspended in fresh medium, plated in new T75 flasks, and cultured for 1 hour. The cell suspensions were centrifuged and resuspended in a growth medium with 5 ng/ml recombinant human basic fibroblast growth factor (bFGF; Thermo Scientific). The muscle satellite cells were cultured at 37 °C under 5% CO_2_ and sub-cultured when they reached 70% confluency. The myoblasts were obtained in this stage.

For cell differentiation, when the myoblasts reached 90% confluence, they were cultured in a differentiation medium composed of DMEM high glucose medium (Thermo Scientific) supplemented with 2% FBS (Thermo Scientific) and 1% antibiotic-antimycotic (Thermo Scientific), at 41 °C under 5% CO_2_, to induce the formation of myotubes and myofibers.

### 2.4. Differentiation of chicken mesenchymal stem cells to adipocyte-like cells

Chicken MSCs were seeded at 6 × 10³ cells/cm² in T75 flasks and 24-well plates with DMEM high glucose medium (Thermo Scientific) supplemented with 10% FBS (Thermo Scientific) and 1% antibiotic-antimycotic (Thermo Scientific), at 39 °C under 5% CO_2_. When the cells reached 60% confluence, the medium was changed to induce transdifferentiation. The transdifferentiation medium was composed of DMEM/F12 containing 12 μg/ml of soy lecithin (L-α-Phosphatidylcholine). After 7 days the medium was supplemented with 10 μg/ml of insulin (Invitrogen). The transdifferentiation medium was replaced every 2 days for 21 days.

### 2.5. Microscopy of lipid accumulation

The differentiated cells were fixed with 4% paraformaldehyde for 30 minutes and washed three times with DPBS on day 21. Lipid staining was performed using two methods: HCS LipidTOX Red Neutral Lipid Stain (Thermo Scientific), diluted 1:1000 according to the manufacturer’s protocol, and Nile Red staining, prepared by diluting the stock solution (1 mg/mL) to a working concentration of 0,5 µg/mL in DPBS. For both staining methods, the cells were counterstained with Hoechst 33342 (Thermo Scientific) at 1 µg/mL in DPBS for 10 minutes at room temperature. After staining, the cells were washed with DPBS and imaged under fluorescence light microscopy (EVOS M7000 Imaging System, Thermo Scientific).

### 2.6. Immunofluorescence staining and imaging

Chicken myoblasts and myotubes were fixed with 4% paraformaldehyde for 20 minutes at room temperature and washed three times with 0.1% Tween-20 in PBS. Permeabilized with 0.5% Triton X-100 for 15 minutes and washed three times with 0.1% Tween-20 in PBS. Cells were blocked with 5% goat serum or 3% bovine serum albumin (BSA) for 1 hour at room temperature and washed three times with 0.1% Tween-20 in PBS. Subsequently, the primary antibodies were added separately: mouse monoclonal anti-Pax7 conjugated Alexa fluor 488 (Santa Cruz Biotechnology) diluted 1:50, mouse monoclonal anti-Myogenin (F5D, Santa Cruz Biotechnology) diluted 1:50, mouse monoclonal anti-Desmin (D33, Thermo Scientific) diluted 1:50, mouse monoclonal anti-MyoD (5.8A, Thermo Scientific) diluted 1:100, mouse monoclonal anti-Myosin 4 (MF20, Thermo Scientific) diluted 1:100, rabbit polyclonal anti-Myf5 (Abcam) diluted 1:100, and rabbit polyclonal anti-ITGA7 (Sunlong Medical) diluted 1:100 in 1% BSA and 0.1% sodium azide, and incubated overnight at 4 °C. Thereafter, the cells were washed three times with 0.1% Tween-20 in PBS, incubated with the secondary antibodies goat anti-mouse IgG or goat anti-rabbit IgG conjugated Alexa Fluor 488 (Thermo Scientific) diluted 1:800 in 1% BSA and 0.1% sodium azide, for 1 hour at 37 °C, and washed three times with 0.1% Tween-20 in PBS. F-actin was counterstained with Rhodamine Phalloidin (Thermo Scientific) diluted 1:500 in DPBS for 30 minutes at room temperature, and nuclei were counterstained with Hoechst 33342 (Thermo Scientific) diluted to 1 µg/ml in DPBS for 10 minutes at room temperature, and washed three times with 0.1% Tween-20 in DPBS. Stained cells were visualized under fluorescence light microscopy (EVOS M7000 Imaging System, Thermo Scientific).

### 2.7. Quantitative RT-PCR

For genetic characterization, total RNA from chicken cells was extracted using TRIzol (Invitrogen) associated with the RNeasy Mini Kit (Qiagen), according to the manufacturer’s recommendations. DNA digestion was performed on the column using RNase-Free DNase Set (Qiagen). RNA samples were quantified using the NanoDrop 2000 spectrophotometer (Thermo Scientific). Complementary DNA (cDNA) was synthesized using the SuperScript III First-Strand Synthesis SuperMix (Invitrogen), according to the manufacturer’s instructions. RT-qPCR reactions were performed using the QuantiNova SYBR Green PCR Kit (Qiagen), with concentration adjustments of each primer set.

In chicken embryonic stem cells (blastoderm), mesenchymal stem cells and myoblasts, the pluripotency genes were evaluated: chicken (c) *cOCT4*, *cSOX3*, *cNANOG*, *cSALL4* and *cCLDN3*, using primer sets described by Giotis et al. [15]; and *cKIT* and *cLIN28A*, described by Han et al. [16]. In dedifferentiated adipocytes, the genes *cPPARG*, *cADIPOQ*, *cPCK1*, *cADRP*, and *cFABP4* were evaluated, using primer sets described by Pasitka et al. [13]. In chicken myoblasts and myotubes, the following genes were evaluated: *cPAX7* and *cMYOD*, using primer sets described by Hong and Do [17] *cMYMK* and *cMYH1E*, described by Ju et al. [18]. In myoblasts and myotubes, the extracellular matrix genes were evaluated: *cCollagen I* α*1*, *cCollagen I* α*2*, *cLaminin*, *cFibronectin*, and *cElastin*, using primers sets described by Ma et al. [19].

Verification of chicken species DNA was performed by RT-qPCR for the *MT-CYB Gallus gallus* gene, described by Pasitka et al. [13], in primary chicken cells (dedifferentiated adipocytes, myoblasts, and myotubes).

The runs were executed on an ABI 7500 Real-Time PCR System (Applied Biosystems), and each sample was amplified in triplicate using 50 ng of cDNA. Relative gene expression was calculated using the formula 2^-ΔCt^ after normalization with the reference gene *cTBP*.

### 2.8. Nutritional analysis

Total protein content in chicken myoblasts was quantified via Dumas method, using Leco FP-528 (St. Joseph, Michigan, USA) equipment, following the AOAC Official Method 992.15. To protein determination, 0.2 g (± 0.0001) of cells were weighted in a tin (Sn) crucible, then placed in the autosampler carousel for further decomposition at 850 °C at O2 atmosphere. Nitrogen content is determined by external calibration with an analytical calibration curve prepared with EDTA (Leco calibration sample P/N 502/092). The nitrogen content was subsequently converted to protein content using an appropriate nitrogen-to-protein conversion factor, ensuring accurate and reliable results.

### 2.9. Biomass production

To obtain muscle biomass, chicken myoblasts were cultured in T75 flasks with DMEM high glucose (Thermo Scientific) supplemented with 10% FBS (Thermo Scientific) until reaching 60% confluence. Cell fusion was induced by replacing the medium with DMEM high glucose (Thermo Scientific) containing 2% FBS (Thermo Scientific), promoting myotube formation. The resulting myofibers were mechanically processed and incubated overnight at 39 °C with a 15% transglutaminase solution.

Adipogenic biomass derived from MSCs differentiated into adipocytes medium supplemented with 12 µg/mL soy lecithin and 10 µg/mL insulin. After differentiation, cells were collected, washed 2 times with DPBS (Thermo Scientific), and incubated with transglutaminase under conditions similar to those applied to the muscle biomass.

### 2.10. Chicken myoblasts cultivation on commercial microcarriers

To evaluate the adhesion of chicken myoblasts to commercial microcarriers as a preliminary step for potential cell culture scale-up, we selected microcarriers provided by Cellva Ingredients.

Microcarriers were prepared according to the manufacturer’s instructions. Initially, the storage solution was removed, and the microcarriers were washed in DPBS (Thermo Scientific). This washing step was repeated to ensure thorough cleaning. The DPBS (Thermo Scientific) was removed, and the microcarriers were transferred to a sterile plate. Subsequently, the microcarriers were equilibrated in the culture medium by adding 1 mL of medium per gram of microcarriers.

Chicken myoblasts were previously cultured in T75 flasks using DMEM high glucose (Thermo Scientific) supplemented with 10% FBS (Thermo Scientific), 100 U/mL penicillin, and streptomycin (Thermo Scientific), under a humidified atmosphere of 5% CO_2_ at 39 °C. Upon reaching confluency, the cells were detached using 0.25% trypsin-EDTA (Thermo Scientific), resuspended in a medium, and then used for the experiments.

The myoblasts were then seeded onto the microcarriers at a density of 3 x 10L cells per gram of microcarriers. The cell suspension was incubated with the microcarriers in a small volume of medium for at least 3 hours to allow for initial cell adhesion to the material. After this period, an additional medium was added, and the culture was maintained for 4 days, with medium changes every two days. The morphology of the myoblasts on the microcarriers was observed under light microscopy (EVOS M7000 Imaging System, Thermo Scientific).

## 3. Results

### 3.1. Chicken mesenchymal stem cells and embryonic stem cells proliferate stably

The culture of chicken stem cells was stable, with the ability to self-renew and differentiate into different cell types, such as the transdifferentiation into adipocytes demonstrated here.

In the genetic characterization of primary cells isolated from chicken embryos, the expression of pluripotency genes varied among the different types of chicken cells. Embryonic stem cells (blastoderm), which represent an early stage of embryonic development, exhibited high expression of the evaluated genes, reflecting their pluripotency. In contrast, mesenchymal stem cells, which possess the capacity for self-renewal and differentiation, showed moderate expression, demonstrating their restricted pluripotency. Myoblasts, which are muscle precursor cells committed to differentiating into muscle fibers, exhibited low or undetectable expression of these genes, indicating a loss of pluripotency and functional specialization (Figure 1D).

### 3.2. Myogenic differentiation of muscle satellite cells produce myotubes

Chicken myoblasts were derived from the primary culture of chicken embryo muscle tissue. The established myoblasts presented typical myoblast morphology, which is a fibroblast-like shape with a slightly smaller size (Figure 1A). The cells elongated after reaching confluence and being cultured in a differentiation medium. Cell elongation is a sign of myogenesis, which is the result of fusion between myoblasts, forming linear and multinucleated myotubes (Figure 1B; 1C). Culturing the myoblasts in differentiation medium activated transcription factors, promoting the fusion of several precursor cells to form myotubes, which subsequently developed into myofibers (Sup. Video 1).

To confirm that these established cells were myoblasts and myotubes, they were further characterized by immunocytochemical analysis using antibodies against paired box 7 (PAX7), myogenic factor 5 (MYF5), myogenic determination (MYOD), integrin alpha 7 (ITGA7), myogenin (MYOG), myosin heavy chain (MYHC), and desmin (DES), both in chicken myoblasts (Figure 2A) and chicken myotubes (Figure 2B). The myotubes exhibited more positive staining for MYOG, MYHC, and DESMIN, demonstrating that the myoblasts differentiated into linear and multinucleated myotubes.

The expression of the evaluated genes in chicken myoblasts and myotubes regulates the processes of proliferation, differentiation, and cell fusion that form muscle tissue. The *cPAX7* and *cMYOD* genes were more highly expressed in myoblasts, and the expression of these genes decreased in myotubes (Figure 1E). Their expression decreased substantially during differentiation, indicating a transition to a more differentiated phenotype. The *cMYMK* and *cMYH1E* genes had low expression in myoblasts, and their expression increased in myotubes (Figure 1E). Their increased expression is essential for cell fusion and is consistent with their association with muscle fiber maturation.

The extracellular matrix (ECM) is essential for the development, organization, and functionality of these cells. Chicken myoblasts and myotubes play a central role, as they are the precursors of muscle fibers. The genes *cCollagen I* α*1*, *cCollagen I* α*2*, *cLaminin* and *cFibronectin* were highly expressed in myoblasts and myotubes, while the Elastin gene was lowly expressed (Figure 1F). RT-qPCR for the mitochondrially encoded cytochrome B (*MT-CYB*) gene showed that the primary cells isolated and differentiated here originate from the species *Gallus gallus* (data not shown).

### 3.3. Chicken mesenchymal stem cells differentiated into adipocytes

In differentiated adipocytes derived from chicken MSCs, the evaluated genes are associated with the formation and functionality of adipose tissue, which is essential for reproducing the sensory characteristics of meat. In the MSCs morphological changes were observed within four days of induction, with cells adopting a rounded shape characteristic of adipocytes (Figure 3). On day 14, lipid staining using HCS LipidTOX Red Neutral Lipid Stain (Figure 4A; 4B) and Nile Red (Figure 4C; 4D) validated lipid accumulation. Both staining methods revealed the presence of intracellular neutral lipids, which could be triglycerides and cholesterol esters, consistent with the maturation of adipocytes [20].

Gene expression analysis by RT-qPCR shows that *cPPARG* exhibited the highest expression, reflecting its critical role as a master regulator of adipogenesis [21]. Genes such as *cFABP4* and *cADIPOQ*, associated with fatty acid transport and metabolic regulation, were expressed, indicating the acquisition of functional adipocyte properties [22]. The expression of *cPCK1*, involved in lipid and glucose metabolism, validated the successful differentiation process (Figure 4E).

### 3.4. Chicken muscle and adipogenic cells produced biomass

Muscle and adipogenic biomass were produced using established protocols. Approximately 0.3 g of muscle biomass was obtained after culturing chicken myoblasts for 30 days, followed by mechanical harvest and incubation using 15% transglutaminase solution. Similarly, adipogenic biomass, derived from MSCs differentiated into adipocytes, yielded approximately 0.5 g after 14 days of adipogenic induction. Both biomass types were processed into macroscale constructs, demonstrating structural integrity and scalability (Figure 5).

### 3.5. Nutritional analysis revealed myoblasts protein content

The protein content of the chicken myoblast sample was determined using the Dumas combustion method, in accordance with the AOAC Official Method 992.15. Analysis revealed that the cultured myoblasts contained 10.63% total protein on a dry weight basis.

### 3.6. Isolated chicken myoblasts adhered to microcarriers

After 24 hours of incubation, primary chicken myoblasts were observed to adhere efficiently to microcarriers. By 96 hours, an increased density of adhered cells was evident, indicating not only sustained adhesion but also active proliferation over time. These observations suggest that the evaluated microcarriers provide a suitable surface for myoblast attachment and expansion, supporting their potential use in large-scale bioprocesses for cultivated meat production (Figure 6).

## 4. Discussion

To create mimetics of conventional meat using cell culture for protein production from animal cells, the primary goal is to develop mature muscle tissue. Muscle tissue is composed of myofibers (myotubes) – long, multinucleated cells that contract to generate force and enable movement. Besides myofibers, muscle tissue contains other essential cell types, including fibroblasts and adipocytes, all of which contribute significantly to the structural integrity and function of the tissue. Along with this complex cellular environment are ECM proteins that provide structural support, enabling cells to attach, interact, and maintain the cohesive architecture of tissue [23]. Together, these components give skeletal muscle tissue its distinct properties of skeletal muscle tissue in meat. However, it is still challenging to recreate these characteristics in vitro, as in the production of cultured meat. A primary difficulty lies in co-culturing diverse cell types, each with unique characteristics and requirements. In this study, we explored a culture approach by deriving multiple meat components – muscle, and fat – from two cell types, and comparing their yields. Specifically, we utilized chicken embryonic and mesenchymal stem cells to generate muscle tissue, and fat storage in a carefully regulated monolayer culture environment, as well as in three-dimensional cultures using microcarriers. This approach allowed us to cultivate meat with the desired characteristics, proliferating both embryonic and mesenchymal stem cells in the desired quantity and then differentiating them, thus advancing the development of cultured meat with greater similarity to its conventional counterpart.

The analysis of pluripotency gene expression revealed high expression levels in stem cells, pluripotent nature, and their ability to self-renew and differentiate into various cell types. Jean et al. [24] also demonstrated that embryonic cells expressed classical pluripotency-related genes, such as *OCT4*, *NANOG*, *SOX3*, and *SALL4*. Genes such as *OCT4* and *NANOG* play essential roles in maintaining the pluripotent state in chicken ESCs [25], ensuring their ability to remain undifferentiated. MSCs showed moderate expression compared to ESCs, reflecting more restricted pluripotency. MSCs are multipotent stem cells that have the ability to differentiate into osteocytes, chondrocytes, adipocytes, and myocytes [26]. Here, MSCs were transdifferentiated into adipocytes as well as into muscle satellite cells, in which myoblasts differentiated into myotubes.

Skeletal muscle tissue consists not only of mature, multinucleated muscle fibers but also a diverse array of supporting cell types. Muscle satellite cells, as myogenic progenitors, exhibit a robust regenerative capacity, differentiating readily into myotubes and mature myofibrils. Environmental factors, particularly temperature, significantly influence the activity of muscle satellite cells, with effects that can either enhance or inhibit their functionality. In chickens, muscle satellite cell proliferation and differentiation are highly sensitive to changes in temperature [27,28]. In this study, a temperature of 41 °C was applied to support satellite cell differentiation as well as myotube proliferation.

During the differentiation of muscle satellite cells into myofiber, we previously observed a large population of myoblasts cells, characterized by lower or absent expression of these pluripotency genes, reflecting their specialized role in muscle tissue formation. The progression of myoblasts into myofibers is marked by diminished pluripotency and increased expression of muscle-specific genes, underscoring their commitment to a muscle lineage. Our findings demonstrate that the *cPAX7* and *cMYOD* genes are highly expressed in myoblasts but are decreased in myotubes, corroborating previous findings. The reduction in *PAX7* expression signals that progenitor cells have exited the proliferative state and are moving toward differentiation [29]. This is consistent with the role of *MYOD* as a key regulator in the cell cycle transition to myogenic commitment, promoting the expression of differentiation-related genes [30]. In contrast, the *cMYMK* and *cMYH1E* genes exhibit low expression in myoblasts but increase significantly during differentiation into myotubes, underscoring their importance in cell fusion and muscle fiber maturation. *MYH1E*, an essential structural protein in mature muscle fibers, is widely used as a marker of differentiation. Similarly, *MYMK* is essential for the cell fusion process, facilitating the formation of multinucleated myotubes [31]. Identification of cells belonging to the *Gallus gallus* species using RT-qPCR for the *MT-CYB* gene confirms the origin of the cells, ensuring the authenticity of the cultures. These results provide valuable insights into the molecular mechanisms underlying myogenesis and highlight the dynamic expression profiles of myogenic markers during muscle development.

The ECM plays a crucial role in the development, maintenance, and functionality of muscle cells, including both myoblasts and myotubes. It provides structural support, facilitates cell signaling, and organizes tissue architecture. The genes *cCollagen I* α*1*, *cCollagen I* α*2*, *cLaminin* and *cFibronectin* are essential for cell maintenance and differentiation. However, the low expression of *cElastin* reflects the specific microenvironment requirements of muscle tissue [32].

The screening of myoblast and myotube populations resulted in a culture of mononuclear cells with distinct morphology and protein expression patterns compared to muscle satellite cells. A robust immunocytochemistry approach was applied to characterize chicken myoblasts and myotubes, using key markers such as PAX7, MYF5, MYOD, ITGA7, MYOG, MYHC, and DES. These markers are essential in different stages of myogenesis, including proliferation, differentiation, and muscle fiber formation. The myotubes obtained after myogenic differentiation showed strong staining of MYHC, a terminal differentiation marker of skeletal muscle cells. Differentiation into sufficiently mature myofibers created through the fusion of myoblast cells, together with cell proliferation, are important parameters for the production yield and quality of cultured meat. In addition, myoblast-derived biomass showed nutritional viability, with a protein content of 10.63%, supporting its potential as a functional component for cultured meat formulations.

Beyond muscle protein, lipid content is a critical factor in the quality of meat. While muscle cells have a limited capacity to store fat, adipocytes are responsible for generating intramuscular fat, which constitutes approximately 80% of the fat in meat. This fat is critical for imparting juiciness, tenderness, and aroma to meat, with higher fat content enhancing the flavor during cooking [33,34]. Consequently, fat is primarily composed of adipocytes with a high concentration of lipid droplets, which are primarily deposited in fat cells within the tissue [33]. To accurately replicate the sensory characteristics of intramuscular fat in cultured meat, co-culturing muscle and fat cells is essential. For instance, co-culturing preadipocytes with myoblasts can potentially elevate intramuscular fat content, improve tenderness, and enhance flavor intensity in the final product [35]. However, co-culturing diverse cell types presents technical challenges, as each cell type requires a distinct, optimized environment to develop and differentiate effectively. Shared culture conditions may be suboptimal for one cell type, leading to hindered cell growth and efficiency [36]. Research has shown that adipocytes developing in close proximity to muscle cells can modulate myogenesis, thereby influencing the development and characteristics of muscle tissue [37]. To overcome these challenges, identifying the most suitable cell source and optimizing conditions for differentiation into either muscle or fat cells are critical steps in achieving the desired genotypic and phenotypic outcomes for cultured meat.

The adipogenic capacity of preadipocytes can be evaluated through alterations in transcription factor expression and cell cycle characteristics. The initially adhered MSCs were induced to differentiate into preadipocytes. Adipogenesis depends on the essential transcription factor peroxisome proliferator activating receptor gamma (PPAR-γ) [38]. To turn on the adipogenic transcriptional program, PPAR-γ interacts with C/EBP family transcription factors [39]. Together with lipogenic genes including fatty acid synthase (FAS) and fatty acid binding protein 4 (FABP4), mature adipocytes preserve the expression of PPAR-γ, widely regarded as an adipogenic marker. Based on these characteristics, we hypothesized that these cells may be fibro-adipogenic progenitor cells (FAPs) and confirmed their expression of FABP4, as well as other factors previously implicated in adipogenesis, including PPAR-γ and ADIPOQ. These FAPs demonstrated lipid droplet accumulation, as well as strong induction of adipocyte marker genes when treated with a differentiation medium containing adipogenic inducers. As FAPs are a primary source of intramuscular fat depots in vivo, we suggest that MSC-derived cultured fat could more accurately resemble traditional adipose tissue compared to fat produced from other cell types, such as fibroblasts. Notably, the lipid accumulation rate in MSC-derived FAPs was higher than those from broiler ESCs. Furthermore, FAPs can be efficiently co-cultured with muscle satellite cells, enhancing the potential for a viable bioprocess in cultured meat production. To further optimize cell sorting strategies, we conducted an extensive characterization of the immunogenotypic and phenotypic profiles of these FAPs cells.

Our findings demonstrate the feasibility of protocols for differentiating MSCs into adipocytes and producing cellular biomass with desirable characteristics for cultivated meat production. Adipogenic differentiation was validated through gene expression analysis and lipid accumulation, consistent with previous studies emphasizing the roles of genes such as PPARG and FABP4 in adipogenesis and intracellular fatty acid transport [38,39]. Additionally, our use of lecithin as an adipogenic inducer is supported by the work of [13], who demonstrated that phosphatidylcholine, a key component of soy lecithin, activates PPAR-γ in chicken fibroblasts, effectively promoting adipocyte formation. This approach eliminates the need for chemically restrictive inducers or hormones, such as insulin and dexamethasone, making it more suitable for food-grade applications.

The successful adhesion and proliferation of primary chicken myoblasts on microcarriers observed in this study demonstrate their potential as a platform for scalable muscle cell cultivation. After 24 hours of incubation, cells were visibly adhered to the microcarriers, and a substantial increase in cell density was observed at 96 hours, indicating active proliferation. These findings highlight the suitability of the tested microcarriers for dynamic suspension culture systems, such as stirred-tank bioreactors, which are essential for large-scale production of cultured meat. Microcarriers offer a significantly increased surface-to-volume ratio, allowing for higher cell yields in reduced volumes and enhanced process control compared to traditional planar systems [40]. The observed compatibility between the chicken myoblasts and microcarrier surface suggests that microcarriers possess adequate surface chemistry, charge, and topography to support anchorage-dependent cell attachment, spreading, and expansion—critical for preserving cell phenotype and myogenic potential [40–42]. Future studies should investigate the performance of these microcarriers under dynamic culture conditions, particularly focusing on their resistance to shear stress, capacity for bead-to-bead transfer, and detachment efficiency—parameters that are decisive for process scalability and downstream cell recovery [43,44].

In addition to microcarrier-based expansion, the structural consolidation of cellular biomass using transglutaminase further supports the development of 3D tissue-like constructs suitable for food applications. This method, which ensures the structural and sensory integrity of the biomass, is consistent with findings by Yuen et al. [45], who used transglutaminase to consolidate cultivated adipogenic cells, resulting in tissues with mechanical properties comparable to native adipose tissue. In our study, the combined use of mesenchymal stem cell transdifferentiation and transglutaminase stabilization enabled the production of lipid-rich biomass, akin to previous experiments, but with enhanced scalability potential for industrial applications.

In summary, we present a primary cell culture strategy to construct multicomponent tissues by developing myogenic and adipogenic microtissues derived from multipotent cells. For cultured meat production, the use of multipotent cells is particularly advantageous due to their higher proliferation rates, which can enhance scalability and efficiency. Alternatively, the immortalization of primary cells, such as myoblasts and preadipocytes, represents another promising avenue for ensuring long-term cell availability and consistent performance in cultured meat applications.

## Supporting information

chicken myoblasts

## 5. Acknowledgements

The authors acknowledge Franciana A. Volpato for laboratory assistance, and Marina Schmitt for the graphic design. VH, KRDS and MAP have a grant from Funding Agency for Studies and Projects (FINEP).

---

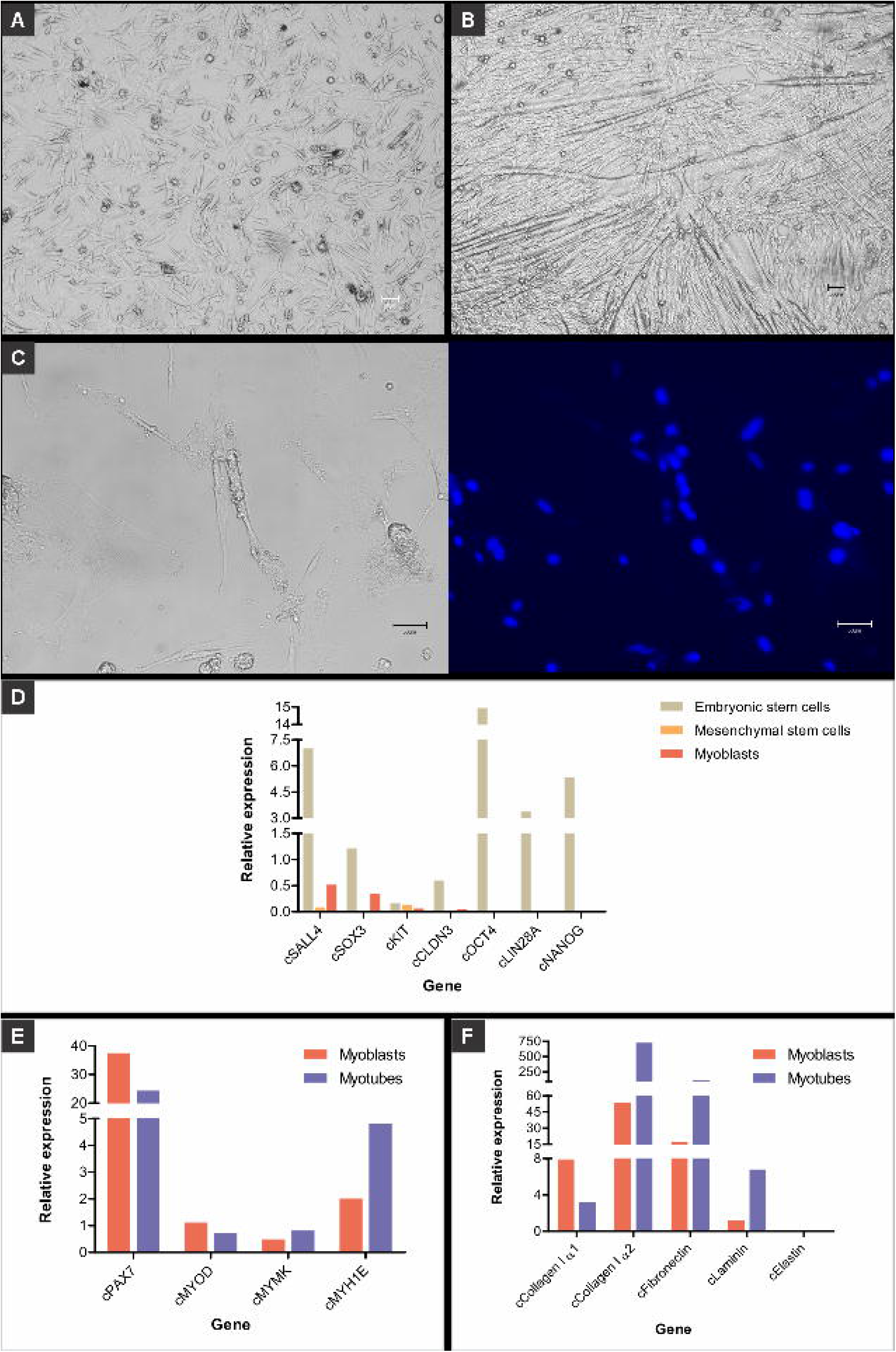

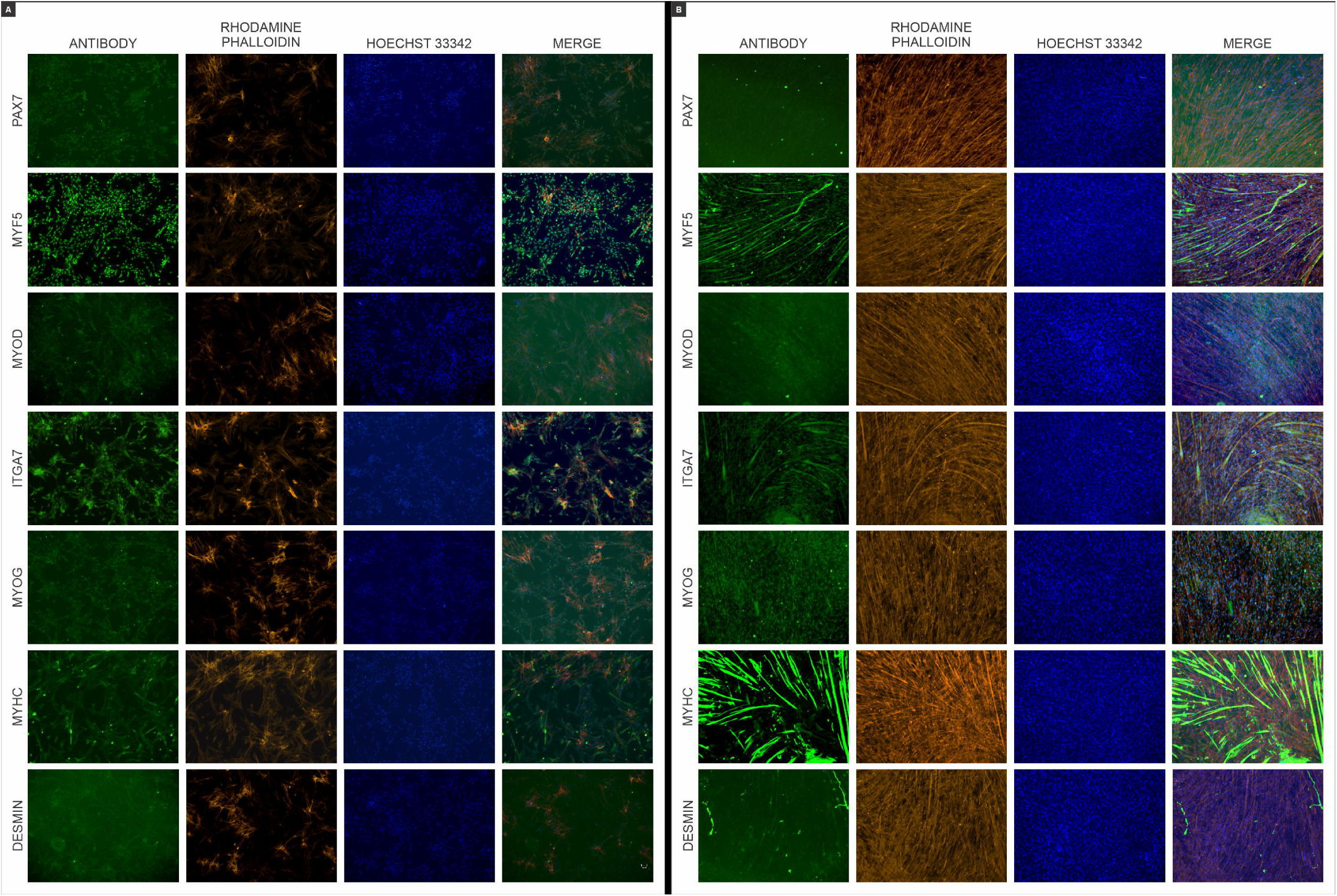

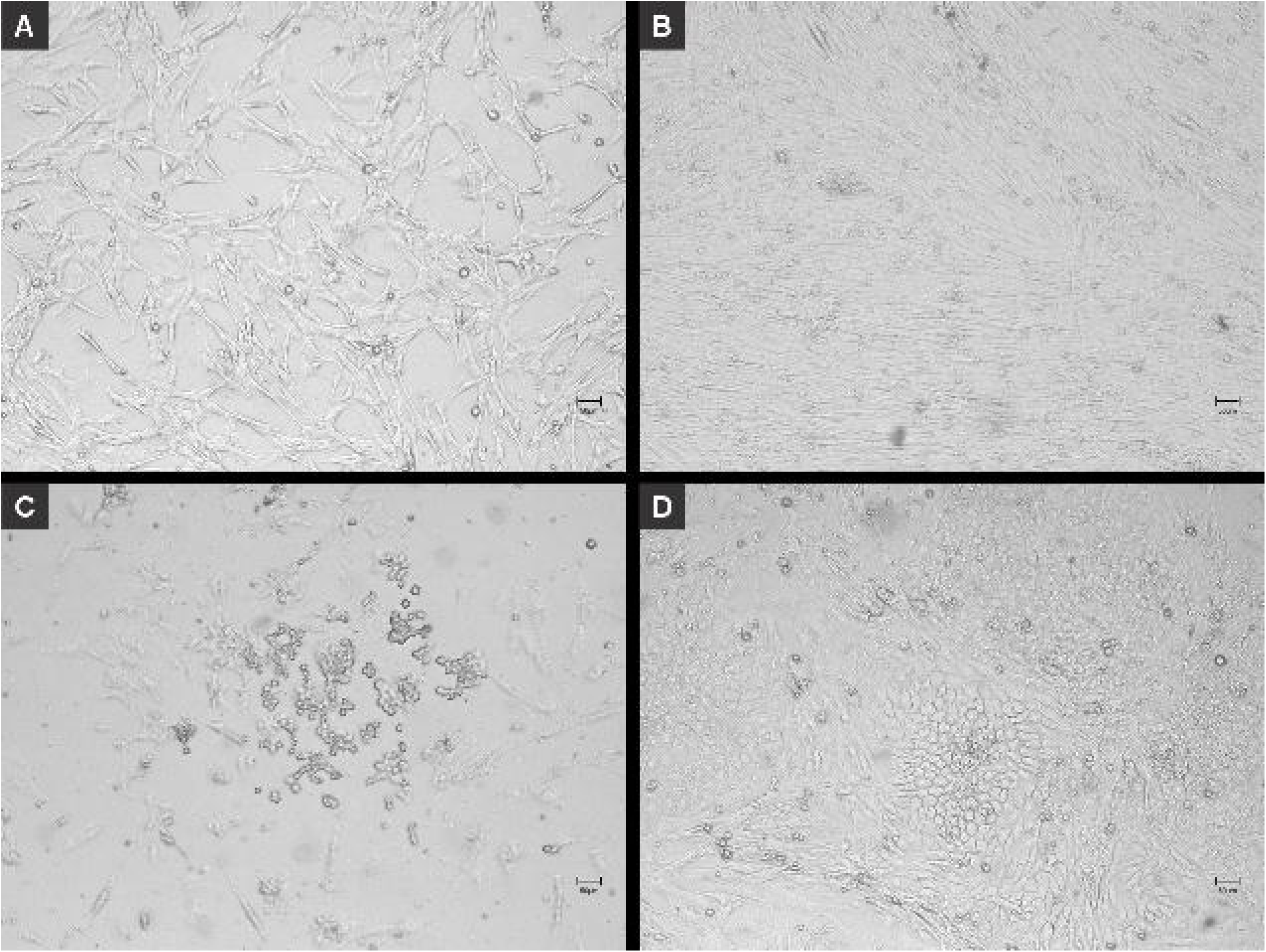

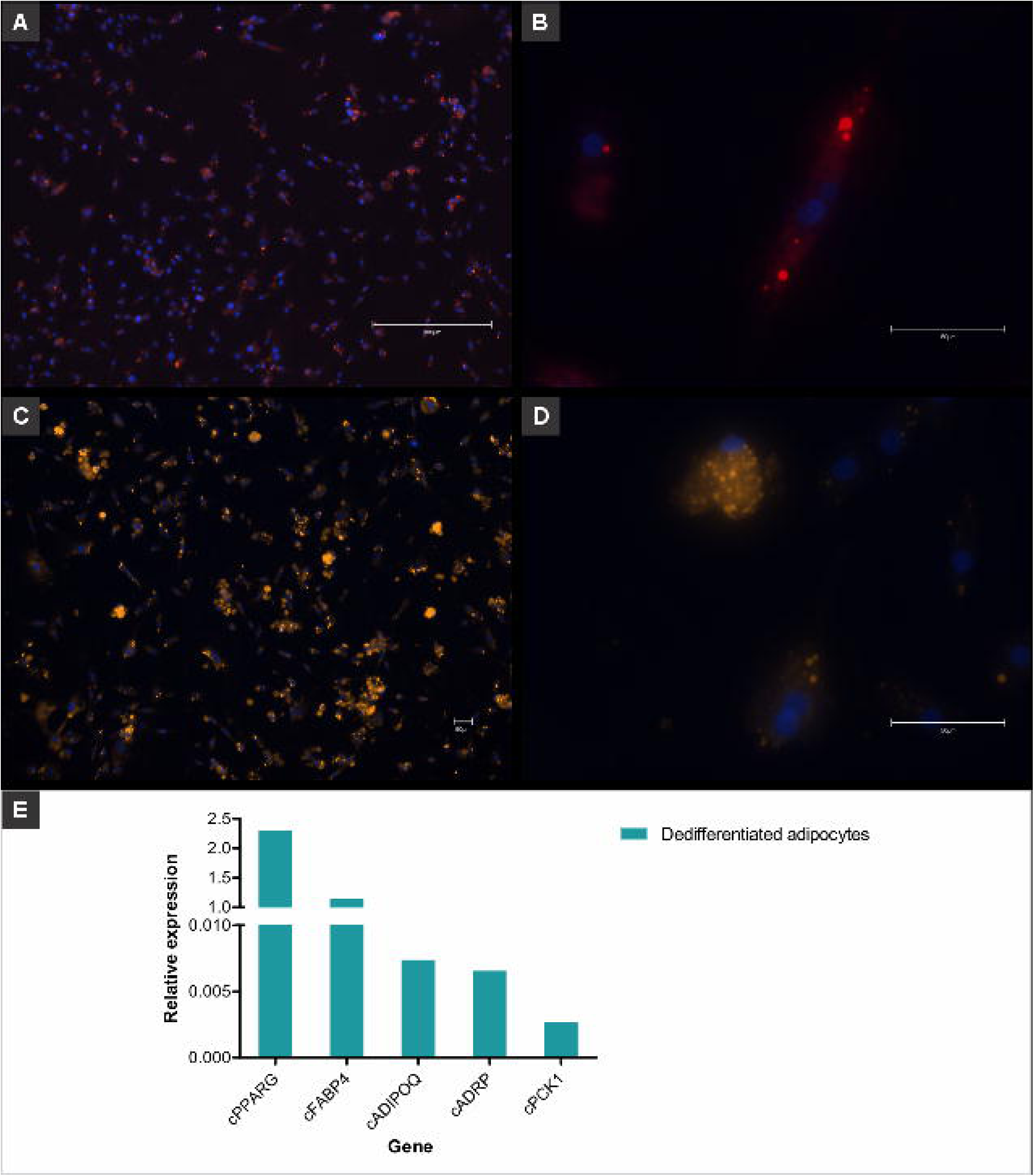

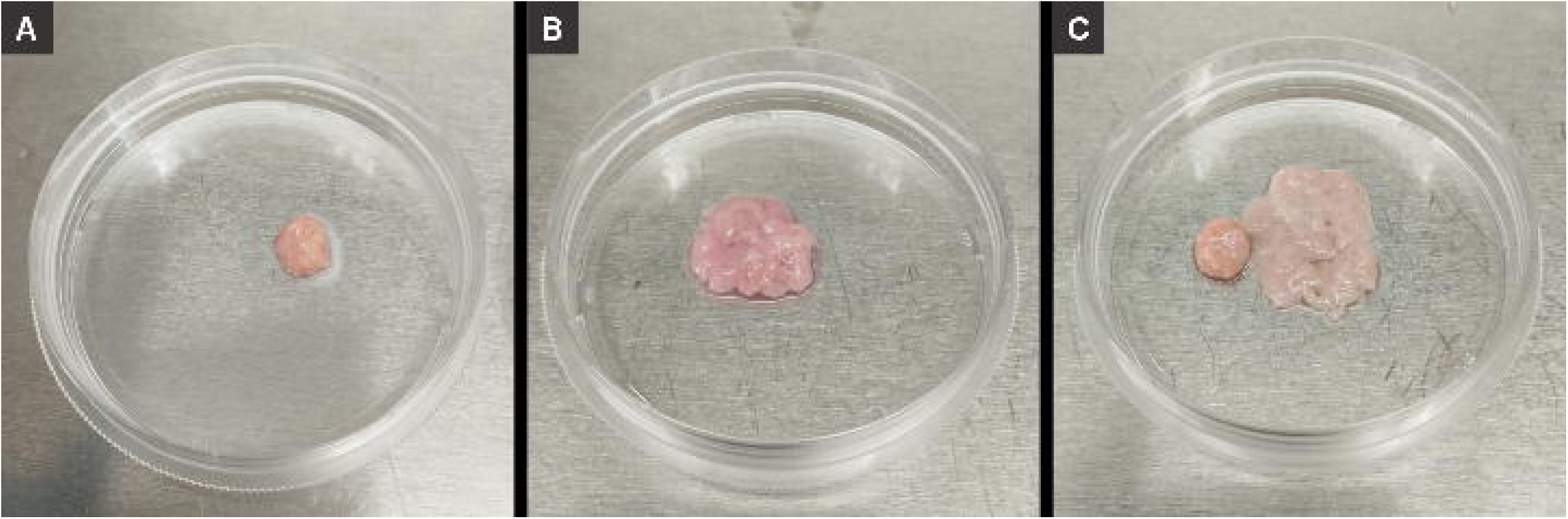

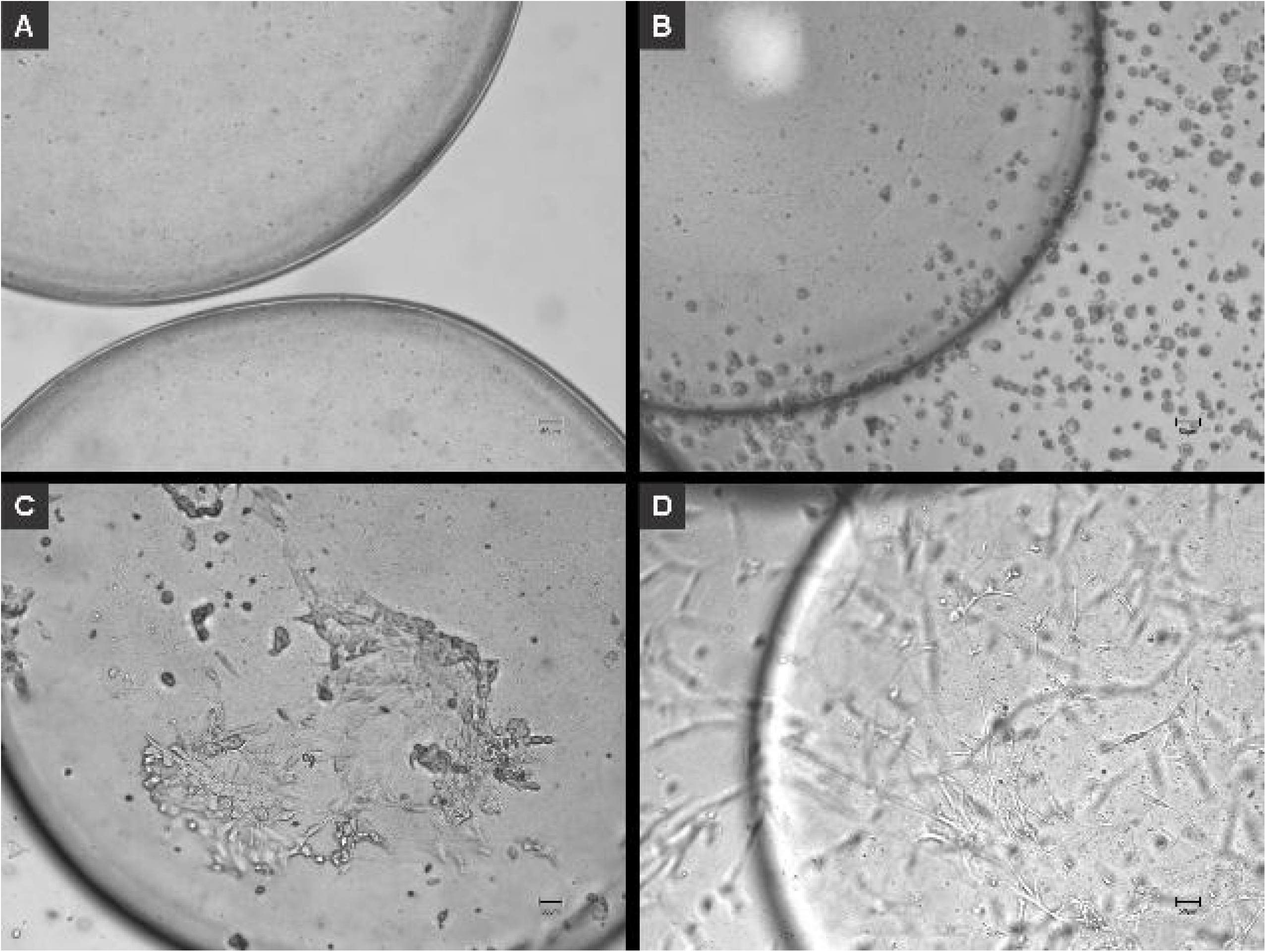

